# RNA polymerase clamp conformational dynamics: long-lived states and modulation by crowding, cations, and nonspecific DNA binding

**DOI:** 10.1101/2020.10.06.327965

**Authors:** Abhishek Mazumder, Anna Wang, Heesoo Uhm, Richard H. Ebright, Achillefs N. Kapanidis

## Abstract

The RNA polymerase (RNAP) clamp, a mobile structural element conserved in RNAP from all domains of life, has been proposed to play critical roles at different stages of transcription. In previous work, we demonstrated using single-molecule Förster resonance energy transfer (smFRET) that RNAP clamp interconvert between three short-lived conformational states (lifetimes ∼ 0.3-0.6 s), that the clamp can be locked into any one of these states by small molecules, and that the clamp stays closed during initial transcription and elongation. Here, we extend these studies to obtain a comprehensive understanding of clamp dynamics under conditions RNAP may encounter in living cells. We find that the RNAP clamp can populate long-lived conformational states (lifetimes >1.0 s) and can switch between these long-lived states and the previously observed short-lived states. In addition, we find that clamp motions are increased in the presence of molecular crowding, are unchanged in the presence of elevated monovalent-cation concentrations, and are reduced in the presence of elevated divalent-cation concentrations. Finally, we find that RNAP bound to non-specific DNA predominantly exhibits a closed clamp conformation. Our results raise the possibility of additional regulatory checkpoints that could affect clamp dynamics and consequently could affect transcription and transcriptional regulation.

## Introduction

Transcription is the first and most highly regulated step in gene expression (*1*). The first step of transcription is the formation of the open complex (RPo), in which the RNA polymerase holoenzyme (RNAP) binds to a promoter DNA fragment and unwinds double-stranded DNA, resulting in a ∼12-14 bp bubble where the individual DNA strands are separated, and single strands of DNA interact with RNAP in a sequence-specific manner (*2*). Extensive structural studies of RNAP, the RPo, and initial transcribing complexes (RPitc) reveal that although RNAP accommodates single strands of DNA in the RNAP active-site cleft, access of double-stranded DNA (width ∼20 Å; (*3*)) inside the cleft is restricted due to the narrow width of the cleft (<20 Å) (*4-7*). This raises the question as to how and when the promoter DNA unwinds on path to the final RPo.

One of the prevailing models in the field implicates the RNAP clamp, a structural element that forms one wall of the active-site cleft, as a crucial player in RPo formation. According to this model, after initial DNA binding, the clamp can swing open, widening the RNAP active-site cleft; this conformational change licenses entry of duplex DNA, subsequent melting of DNA inside the cleft, and clamp closure, resulting in a catalytically competent RPo. Some structural studies have captured the clamp in open clamp conformations that would allow entry of duplex DNA inside the cleft (*8*). Recent structural studies on a σ54 RNAP-promoter complex also captured a conformation where double-stranded DNA is located inside the RNAP cleft with a wide-open clamp conformation (*9*). However, it has not been established conclusively that such conformations are on-pathway to RPo formation. Additionally, motions of the clamp have been proposed to play crucial roles in transcription elongation (*10,11*) and transcription termination (*12*).

It is therefore important to study clamp conformations and conformational dynamics in solution, and in real time, to detect and characterize clamp conformations and conformational dynamics, to determine timescales of the clamp conformational dynamics, to understand the modulation of clamp conformational dynamics by changes in conditions, and to assess the functional relevance of clamp conformational dynamics. Clamp conformation and conformational dynamics can be directly detected and characterized using single-molecule FRET (smFRET), which enables measurement of distances, in the range of 2-10 nm, between pairs of fluorophores incorporated at specific sites within a macromolecule (*13*). One of the major challenges in performing smFRET measurements is the incorporation of a pair of fluorophores at specific sites of interest in the macromolecule of interest. In previous work, we developed a procedure that combines unnatural-amino-acid mutagenesis and Staudinger ligation to enable preparation of RNAP derivatives having the fluorescent probes Cy3B and Alexa 647 incorporated at the tip of the RNAP clamp and the tip of the opposite wall of the RNAP active-site cleft (*13-14*). We first used the resulting doubly labelled RNAP derivatives to monitor clamp conformations in freely diffusing RNAP molecules in solution through smFRET with confocal alternating laser excitation microscopy (confocal ALEX; *13*). We next used the doubly labelled RNAP derivatives to monitor clamp conformational *dynamics* in surface-immobilized RNAP molecules in solution, *in real time*, through smFRET with total internal reflection fluorescence ALEX microscopy (TIRF-ALEX; *15-16*). We observed that, in RNAP holoenzyme, the clamp populates three distinct conformations: an open clamp, a partly closed clamp, and a closed clamp conformation (*13, 15, 16*). We also showed that the clamp switches among these three short-lived conformational states on the millisecond time scale and can be trapped in any one of these three conformations by the RNAP inhibitors myxopyronin, corallopyronin, ripostatin, fidaxomicin (lipiarmycin), and ppGppp (*13, 15, 16*). Additionally, we observed that the clamp adopts exclusively or nearly exclusively, a closed conformation in the RNAP-promoter open complex (RPo), RNAP-promoter initial transcribing complexes (RPitc), and RNAP-DNA elongation complexes (RDe) (*13,15*).

Here, we extend our studies to further our understanding of modulation of clamp conformational dynamics in solution in real time under several physiologically relevant settings. Our results show that the RNAP clamp can populate additional long-lived clamp conformational states and can switch between these states and the previously observed short-lived clamp states. Our results further show that the RNAP clamp exhibits unchanged conformational dynamics in the presence of elevated K^+^, and decreased conformational dynamics in the presence of elevated Mg^2+^; exhibits enhanced conformational dynamics in the presence of molecular crowding; and show that most RNAP molecules engaged in nonspecific interaction with DNA exhibit a closed clamp conformation. Taken together, the results show that clamp conformation and dynamics vary under different solution conditions and offer insight into how modulation of clamp motions might impact transcription regulation.

## MATERIALS AND METHODS

### RNAP preparation

Fluorescently labelled, hexahistidine-tagged *E. coli* RNAP holoenzyme (hereafter “clamp labelled RNAP”) with Cy3B and Alexa647 at positions 284 on the β’ subunit, and 106 on the β subunit, respectively, were prepared using an *in vivo* reconstitution as described in (*16*).

### DNA

Oligonucleotides were purchased from Sigma Aldrich and annealed in hybridization buffer (50 mM Tris-HCl pH 8.0, 500 mM NaCl, 1 mM EDTA). The sequence for the non-specific DNA was generated from the lacCONS promoter sequence (*15*) by full substitution of the −35 and −10 elements, as follows:

5’-AGGCGCTGTCCTTTATGCTTCGGCTCGCCGGTAGTGTGGAATTGTGAGAGCGGATAACAATTTC–3’
3’-TCCGCGACAGGAAATACGAAGCCGAGCGGCCATCACACCTTAACACTCTCGCCTATTGTTAAAG-5’

### Formation of RNAP complexes with non-promoter DNA

RNAP complex with non-specific DNA were prepared by mixing 20 nM of biotinylated-non-specific dsDNA with 50 nM labelled RNAP for 5 min at 37°C in KG7 buffer (40 mM HEPES-NaOH, pH 7.0, 100 mM potassium glutamate, 10 mM MgCl_2_, 1 mM dithiothreitol and 5% glycerol).

### Single-molecule fluorescence imaging of RNAP under different conditions

For all single-molecule experiments except experiments in Figure 3B, a biotin-PEG-passivated glass surface was prepared, functionalized with neutravidin and treated with biotinylated anti-hexahistidine monoclonal antibody (Qiagen) as described (*15*). RNAP immobilisation was performed by adding 100 pM solution of labelled RNAP holoenzyme to the PEGylated surfaces with biotinylated anti-hexahistidine monoclonal antibody (*15*) for 5 min at 22°C in KG7 buffer (40 mM HEPES-NaOH, pH 7.0, 100 mM potassium glutamate, 10 mM MgCl_2_, 1 mM dithiothreitol, 5% glycerol). Observation wells containing immobilised labelled RNAP were washed with 3×50 μl KG7. For experiments performed under different solution conditions the following solution mixtures were added to the observation wells 3 min before recording of the movie:

For experiments in Figure 1: 30 μl KG7 imaging buffer (40 mM HEPES-NaOH, pH 7.0, 100 mM potassium glutamate, 10 mM MgCl_2_, 1 mM dithiothreitol, 5% glycerol and 2 mM TROLOX, plus an oxygen scavenging system consisting of 1 mg/mL glucose oxidase, 40 μg/mL catalase, and 1.4% w/v D-glucose).

**Fig. 1.**
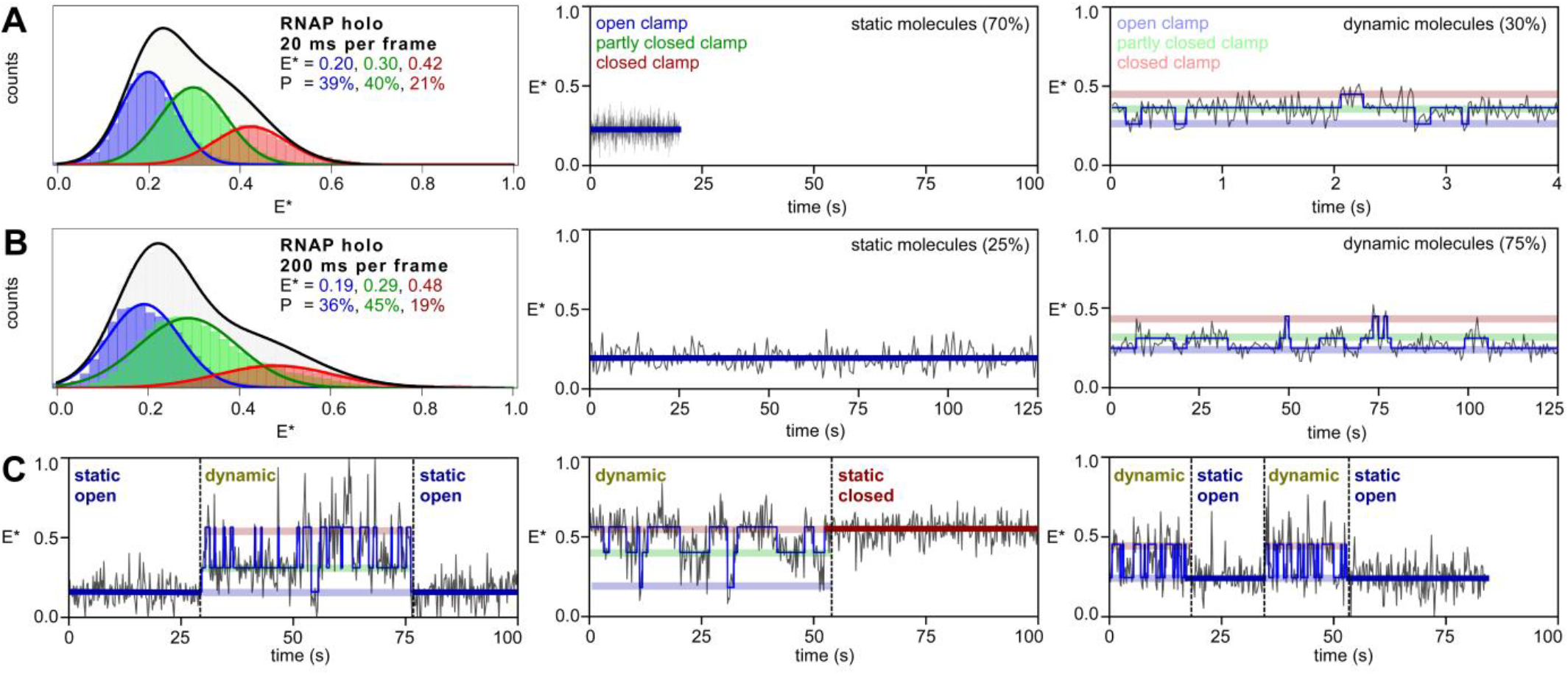
RNAP clamp switches between long-lived and short-lived conformational sub-states. **A**. (*left*) histograms and Gaussian fits of E*, showing HMM-derived E* distributions for open (blue bars), partly closed (green bars) and closed (red bars) clamp states; P, subpopulation percentage. (*middle*) representative time traces of FRET efficiency, E*, showing a molecule in a static open clamp state. (*right*) representative time trace of FRET efficiency, E*, for a dynamic molecule showing hidden-Markov-model (HMM)-assigned states and interstate transitions (blue line). Relative percentages of the populations are inset. Laser powers were 3.5 mW in green and 0.75 mW in red. Frame rate is 20 ms. **B**. (*left*) histograms and Gaussian fits of E*, showing HMM-derived E* distributions (*middle*) representative time trace of FRET efficiency, E*, showing a molecule in a static open clamp state. (*right*) representative time trace of FRET efficiency, E*, for a dynamic molecule showing HMM-assigned states. Laser powers were 0.20 mW in green and 0.075 mW in red. Frame rate is 200 ms. **C**. (*left*) time trace of FRET efficiency, E*, showing transition between a static open clamp status to a dynamic status (*middle*) time trace of FRET efficiency, E*, showing transition between a dynamic status to a static closed clamp status (*right*) time trace of FRET efficiency, E*, showing transitions between dynamic and static clamp status. Laser powers and frame rate as in B.

For experiments in Figure 3A: 30 μl KG7 imaging buffer supplemented with 20% PEG-8000.

For experiments in Figure 2A: 30 μl imaging buffer with 1M KCl (40 mM HEPES-NaOH, pH 7.0, 1000 mM potassium chloride, 10 mM MgCl_2_, 1 mM dithiothreitol, 5% glycerol and 2 mM TROLOX, plus an oxygen scavenging system consisting of 1 mg/mL glucose oxidase, 40 μg/mL catalase, and 1.4% w/v D-glucose).

**Fig. 2.**
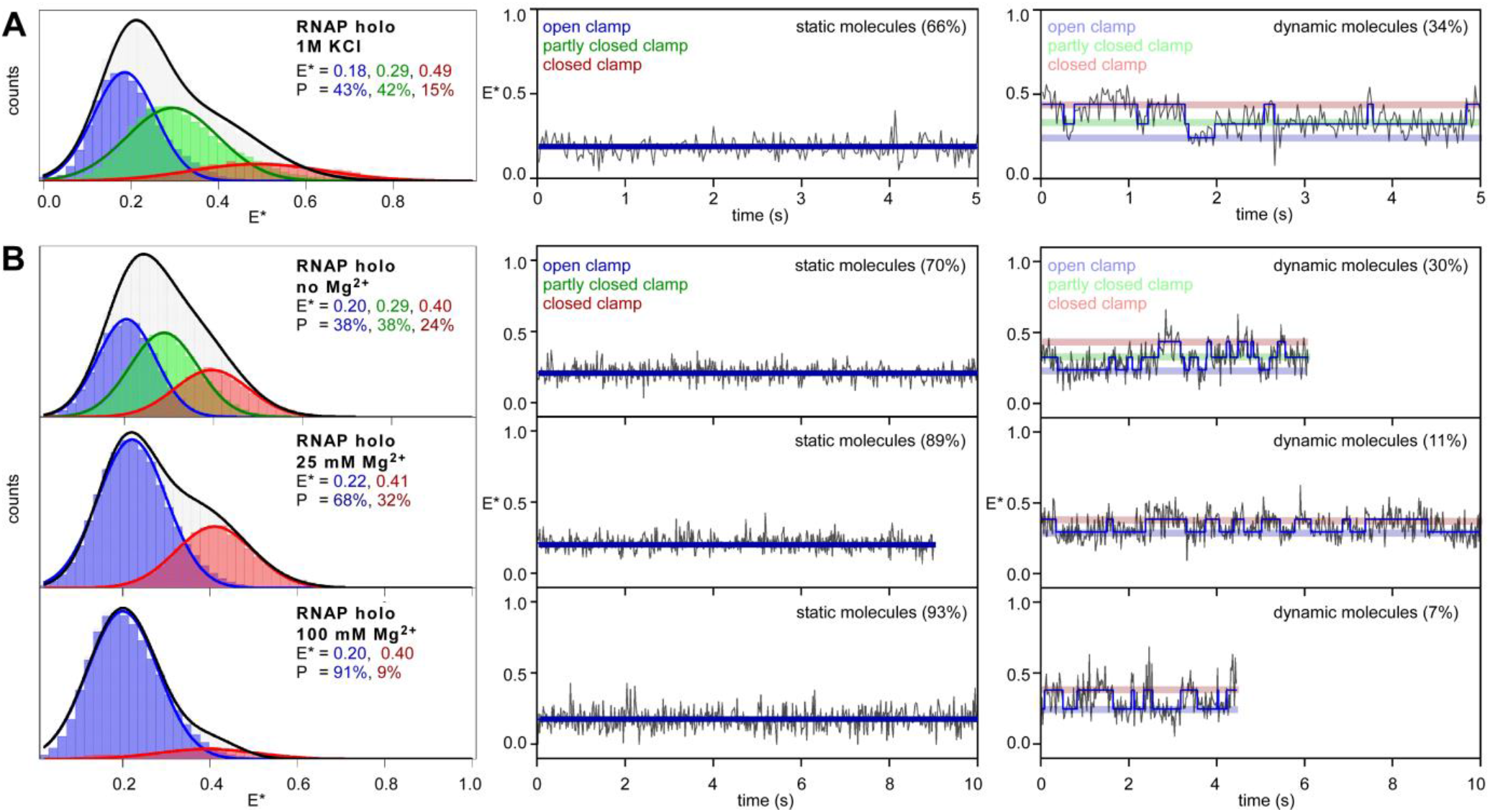
Modulation of RNAP clamp conformation and dynamics in presence of increased concentration of monovalent and divalent salt. **A**. smFRET data monitoring clamp conformation in presence of high K^+^ concentration (1000 mM). (*left*) histograms and Gaussian fits of E*, showing HMM-derived E* distributions; P, subpopulation percentage. (*middle*) representative time trace of FRET efficiency, E*, showing a molecule in a static open clamp state (*right*) representative time trace of FRET efficiency, E*, for a dynamic molecule showing HMM-assigned states, and interstate transitions (blue line). **B**. smFRET data monitoring clamp conformation in presence of increasing Mg^2+^ concentrations (no Mg^2+^, 25 mM Mg^2+^ and 100 mM Mg^2+^). (*left subpanels*) histograms and Gaussian fits of E*, showing HMM-derived E* distributions; P, subpopulation percentage. (*middle subpanels*) representative time traces of FRET efficiency, E*, showing molecules in a static open clamp state (*right subpanels*) representative time trace of FRET efficiency, E*, for a dynamic molecule showing hidden-Markov-model (HMM)-assigned states and interstate transitions (blue line). Relative percentages of the populations are inset. Frame rate was 20 ms and laser powers were 3.5 mW in green and 0.75 mW in red.

For experiments in Figure 2B: 30 μl imaging buffer with no Mg^2+^ or 25 mM Mg^2+^ or 100 mM Mg^2+^ (40 mM HEPES-NaOH, pH 7.0, 1 M potassium chloride, 0 or 25 mM or 100 mM MgCl_2_, 1 mM dithiothreitol, 5% glycerol and 2 mM TROLOX, plus an oxygen scavenging system consisting of 1 mg/mL glucose oxidase, 40 μg/mL catalase, and 1.4% w/v D-glucose).

For experiments in Figure 3B, 100 pM RNAP-non-specific DNA complex was added to a biotin-PEG-passivated glass surface functionalized with neutravidin and incubated for 5 min at 22°C in KG7 buffer. Observation wells containing immobilised labelled RNAP were washed with 3×50 μl KG7 and 30 μl KG7 imaging buffer (40 mM HEPES-NaOH, pH 7.0, 100 mM potassium glutamate, 10 mM MgCl_2_, 1 mM dithiothreitol and 5% glycerol, 2 mM TROLOX, plus an oxygen scavenging system consisting of 1 mg/ml glucose oxidase, 40 μg/ml catalase, 1.4% w/v D-glucose) was added just before movies were recorded.

**Fig. 3.**
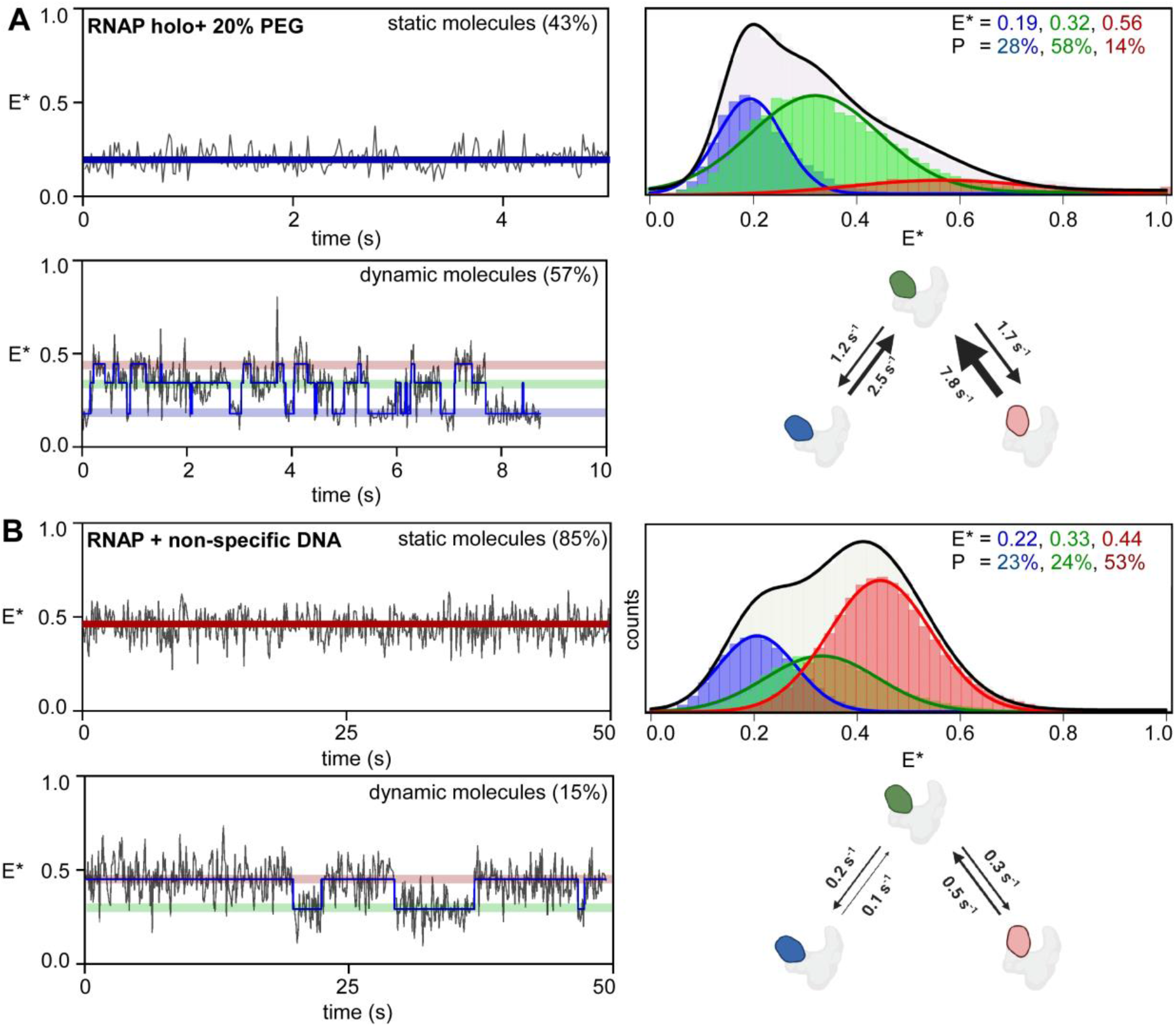
RNAP clamp exhibit enhanced dynamics under conditions of macromolecular crowding and adopts mostly closed conformation when bound to non-specific DNA. **A**. smFRET data monitoring clamp conformation in presence of 20% polyethylene glycol (PEG). (*top left*) representative time trace of FRET efficiency, E*, showing a molecule in a static open clamp state (*bottom left*) representative time trace of FRET efficiency, E*, for a dynamic molecule showing HMM-assigned states, and interstate transitions (blue line). *(top right)* Histograms and Gaussian fits of E*, showing HMM-derived E* distributions; P, subpopulation percentage. Relative percentages of the populations are inset. *(bottom right)* Transition rates between open, partly closed and closed conformations. Frame rate was 20 ms and laser powers were 3.5 mW in green and 0.75 mW in red. **B**. smFRET data monitoring clamp conformaton in RNAP bound to non-specific DNA. (*top left*) representative time trace of FRET efficiency, E*, showing a molecule in a static closed clamp state (*bottom left*) representative time trace of FRET efficiency, E*, for a dynamic molecule showing HMM-assigned states and interstate transitions (blue line). *(top right)* Histograms and Gaussian fits of E*, showing HMM-derived E* distributions; P, subpopulation percentage. Relative percentages of the populations are inset. *(bottom right)* Transition rates between open, partly closed and closed conformations. Frame rate was 100 ms and laser powers were 0.50 mW in green and 0.12 mW in red.

### Single-molecule fluorescence instrumentation

Single-molecule FRET experiments were performed on a custom built objective-type total-internal-reflection fluorescence (TIRF) microscope (*17*). Light from a green laser (532 nm; Samba; Cobolt) and a red laser (635 nm; CUBE 635-30E, Coherent) was combined using a dichroic mirror, coupled into a fiber-optic cable, focused onto the rear focal plane of a 100x oil-immersion objective (numerical aperture 1.4; Olympus), and displaced off the optical axis such that the incident angle at the oil-glass interface exceeds the critical angle, creating an evanescent wave. Alternating-laser excitation (ALEX) was implemented by directly modulating the two lasers using an acousto-optical modulator (1205C, Isomet). Fluorescence emission was collected from the objective, separated from excitation light by a dichroic mirror (545 nm/650 nm, Semrock) and emission filters (545 nm LP, Chroma; and 633/25 nm notch filter, Semrock), focused on a slit to crop the image, and then spectrally separated (using a dichroic mirror; 630 nm DLRP, Omega) into donor and emission channels focused side-by-side onto an electron-multiplying charge-coupled device camera (EMCCD; iXon 897; Andor Technology). A motorized x/y-scanning stage with continuous reflective-interface feedback focus (MS-2000; ASI) was used to control the sample position relative to the objective.

### Image analysis and data processing

Movies of surface-immobilized labeled RNAP were analyzed using the home-built software TwoTone-ALEX (*17*), and the background-corrected intensity-vs.-time traces for donor emission intensity upon donor excitation (I_DD_), acceptor emission intensity upon donor excitation (I_DA_), and acceptor emission intensity upon acceptor excitation (I_AA_) were extracted as described (*17*). For each dataset, we manually inspected intensity time traces and exclude traces exhibiting multiple-step donor or acceptor photobleaching; traces exhibiting donor or acceptor photobleaching in frames 1-50; and traces exhibiting donor or acceptor blinking. The set of selected intensity time traces were used to calculate time traces of apparent donor-acceptor FRET efficiency (E*) and donor-acceptor stoichiometry (S), as described (*18*):

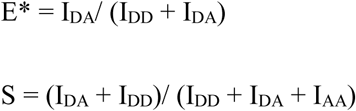

The E* time traces include only data points preceding any donor or acceptor photobleaching events. Two-dimensional E*-S plots were constructed using all selected data points to distinguish species containing donor only (D-only), acceptor only (A-only), and both donor and acceptor (D-A). For species containing both donor and acceptor (D-A), one-dimensional E* histograms were plotted.

### HMM analysis of FRET time-traces

The set of E* time traces selected for analysis were analysed using Hidden Markov Modelling as implemented in the ebFRET software (*19*); and fitted to models with three distinct E* states (the basis for selection of a three-state model for describing clamp conformational dynamics has been extensively discussed in our previous work; Ref. *15*). For experiments in Figure 2B (datasets corresponding to 25 mM or 100 mM Mg^2+^) we estimated that a two-state model fits the data better than a three-state model.

The apparent FRET efficiencies (E*) from the fit to a two- or three-state model were extracted, plotted in Origin (Origin Lab) for the overall population, as well as for the static subpopulation alone, and the dynamic subpopulation alone. The resulting E* histograms linked to each state were fitted to single Gaussian functions in Origin. The resulting histograms provide the equilibrium population distributions of states with distinct E*, define numbers of subpopulations with distinct E*, and, for each subpopulation, define mean E*.

For transition-rate measurements, an HMM analysis was applied to time-traces exhibiting dynamic behaviour. To identify dynamic traces from the set of selected molecules, individual traces were manually checked and those showing anti-correlated changes in the DD and DA channel were identified. Molecules showing greater than three transitions were assigned as dynamic. The data were fitted to a two- or three-state model, and rates of transitions between the states, k_ij_ (where i and j are the states before and after transition), were calculated by multiplying the transition probabilities per frame with the number of frames per second. Mean lifetime in a clamp conformational state, *i*, was estimated from the transition rates out of that state as 1/(∑k_ij_).

### Simulation of FRET traces

We used Hidden Markov Models for our three-state system simulation. The transition probability matrix of the three-state model was calculated from the rates obtained in the short-frame time experiments (20 ms per frame). The matrix was used to generate three state FRET time traces. The time interval was same as the short-frame time (20 ms). To emulate FRET time traces under a long-frame time (200 ms) conditions using the rates obtained in the short-frame time experiments, we binned the short-frame time traces into the long-frame time traces by averaging 10 frames. To estimate a Gaussian noise magnitude in FRET efficiency, we measured the standard deviations of FRET efficiency in static traces from experiments performed at 200ms frame time. The mean of these, 0.068, was then applied to all simulated FRET time traces as the standard deviation of Gaussian noise. The time length of each trace was 600 frames. We generated 200 simulated traces for each condition. This size of the data is selected from the typical size of the experimental data. ebFRET was used to determine transition rates from the traces. All simulations were implemented in MATLAB.

## Results

### Detection of RNAP clamp conformation and conformational dynamics

Consistent with previous smFRET studies on RNAP clamp conformation and conformational dynamics (*13, 15, 16*), the current experiments with surface-immobilised, doubly labelled RNAP derivatives having fluorophores incorporated at the tip of the RNAP clamp and the tip of the opposite wall of the RNAP active-site cleft (“clamp labelled RNAP”) showed that, in RNAP holoenzyme, the RNAP clamp populates three conformational states: (1) open (mean E* ∼ 0.20; 39%), (2) partly closed (mean E* ∼ 0.30; 40%), and (3) closed (mean E* ∼ 0.42; 21%) clamp states (Figure 1A, *left*). Also consistent with previous smFRET studies on RNAP clamp conformation and conformational dynamics (*15, 16*), 30% of the molecules exhibited a “dynamic” clamp conformational status (i.e., exhibited transitions between the three conformational states; Figure 1A, *middle*), and ∼70% of molecules showed “static” behaviour (i.e., did not exhibit detectable transitions between the three conformational states with our observation time of ∼10 s and our observation-time resolution of 20 ms). An inspection of “dynamic” molecules revealed that the RNAP clamp populates short-lived clamp conformational states with average dwells of ∼0.3-0.6 s per state. An estimation of transition rates from these molecules showed that the rates for the three possible clamp-opening transitions, k_closed-partly_closed_, k_partly_closed-open_, and k_closed-open_ were ∼ 3.3 s^-1^, 0.4 s^-1^ and 0.5 s^-1^, respectively, and showed that the rates of the three possible clamp-closing transitions, k_partly_closed-closed_, k_open-partly_closed_ and k_open-closed_, were ∼ 1.1 s^-1^, 1.2 s^-1^ and 0.5 s^-1^, respectively (Fig. S1, *left panel*; Table 1).

**Table 1.** Transition rates between clamp conformational states under different conditions.

| | $K^+$ | $Mg^{2+}$ | $k_{open-partly\ closed}$ | $k_{open-closed}$ | $k_{partly\ closed-open}$ | $k_{partly\ closed-closed}$ | $k_{closed-open}$ | $k_{closed-partly\ closed}$ |
| --- | --- | --- | --- | --- | --- | --- | --- | --- |
| | (mM) | (mM) | ($s^{-1}$) | ($s^{-1}$) | ($s^{-1}$) | ($s^{-1}$) | ($s^{-1}$) | ($s^{-1}$) |
| RNAP | 100 | 10 | 1.18 | 0.55 | 0.42 | 1.14 | 0.50 | 3.35 |
| RNAP | 1000 | 10 | 1.80 | 0.06 | 1.00 | 2.24 | 0.06 | 6.00 |
| RNAP | 100 | 0 | 0.71 | 0.05 | 0.68 | 0.68 | 0.15 | 2.76 |
| RNAP | 100 | 25 | - | 3.27 | - | - | 5.57 | - |
| RNAP +<br>non-specific DNA | 100 | 10 | 0.08 | - | 0.25 | 0.28 | - | 0.47 |
| RNAP +<br>20% PEG-8000 | 100 | 10 | 2.49 | - | 1.23 | 1.75 | - | 7.76 |

### The RNAP clamp populates long-lived dynamic conformational states

Previous work showed that the RNAP clamp adopts exclusively a closed conformation upon formation of RPo (*13, 15*), implying that that all molecules of RNAP holoenzyme that exhibit “static” open and partly closed conformations are able to adopt a closed conformation upon addition of promoter DNA and formation of RPo (*13, 15*). The existence of RNAP molecules that do not show detectable transitions between different clamp states in RNAP holoenzyme, but that, nevertheless, are able to undergo transitions to a closed-clamp conformational state upon formation of RPo, raises the possibility that, in RNAP holoenzyme, the RNAP clamp may populate not only the three characterized sub-second-timescale dynamic conformational states, but also longer-lived dynamic conformational states for which conformational transitions to other states occur on time scales substantially longer than the ∼10 s observation span of the experiments in Fig. 1A and in previous work (*15*).

To assess this possibility, we extended the mean observation span from ∼10 s to ∼120 s. To do this, we increased the frame time from 20 ms to 200 ms, and, to mitigate increased photobleaching with increased observation time, we decreased the laser power by ∼20-fold and observed an increase in observation span from ∼10 s to ∼120 s. We observed that increasing the observation-span from ∼10 s to ∼120 s increased the fraction of molecules of RNAP holoenzyme that show dynamics from ∼30% to ∼75% (Fig. 1A vs. Fig. 1B, *right*), with only few molecules (25%) showing no dynamic behaviour even during the extended observation span (Figure 1B, *middle*). Estimated transition rates from the “dynamic” molecules in the dataset indicated that rates for clamp-opening motions, k_closed-partly_closed_, k_partly_closed-open_, and k_closed-open_, were ∼ 0.5 s^-1^, 0.2 s^-1^ and 0.03 s^-1^, respectively, and that rates of clamp-closing motions, k_partly_closed-closed_, k_open-partly_closed_, and k_open-closed_, were ∼ 0.3 s^-1^, 0.3 s^-1^ and 0.02 s^-1^, respectively (Fig. S1). The observed transition rates are significantly slower, ∼0.02-0.5 s^-1^, as compared to those observed in the previous, short-span experiments (rates of ∼0.5-3.3 s^-1^). The longer-span experiments also showed longer average dwells in individual clamp conformational states of ∼1.5-3.0 s as compared to the shorter-span experiments (dwell times of 0.1-1.0 s).

An inspection of individual FRET time-trajectories showed molecules (∼12%) that populate a very long-lived clamp conformational state (stable for >20 s) before switching to another clamp conformation (Figure 1C, *left*) as well as molecules that exhibit a “dynamic” clamp status (rapid switching between relatively short-lived clamp conformational states) but transition to a very long-lived clamp conformational state at a later time-point (Figure 1C, *middle*). Further, a few molecules (∼2%) show repeated switching between short-lived clamp conformational states and a very long-lived clamp conformational state (Figure 1C, *right*). The observed interconversions between very long-lived and short-lived conformational states, and the overall decreased rates for transitions among different clamp conformations, may be a result of two possible scenarios. In the first scenario, we may be missing transitions between clamp conformations as a result of slower frame rate of the experiment, i.e., entire dwells corresponding to different clamp conformational states are being missed, as the dwell times in these states are shorter than the time of individual frames (i.e., dwells shorter than 200 ms), resulting in artificially long dwells in a clamp conformational state, which would appear as long-lived conformational states and lead to an overall lowering of observed transition rates. From the transition rates measured from the 20-ms frame experiments, we estimate that ∼30%, 41% and 57% dwells in the open, partly closed and closed clamp states would be missed in experiments with 200-ms frames. In the second scenario, there may be hidden transitions between kinetically different clamp conformational states with similar FRET values (e.g., a long-lived and short-lived clamp conformational state with same FRET values), resulting in long dwells. As an example, transitions between a long-lived open clamp state and a short-lived open clamp state would result in an artificially longer dwell in the open-clamp state where all transitions between these two states are missed as they have identical FRET values leading to long dwells in the open-clamp conformational state. It is possible that such dwells in dynamic molecules are mostly missed in experiments with shorter observation span (∼10 s) while we are able to capture a significant proportion of such dwells in dynamic molecules from experiments conducted with extended observation span (∼120 ms).

To understand the origin of this behaviour, we simulated FRET time trajectories at a frame rate of 200-ms per frame using transition rates obtained from the 20-ms frame time experiments in Fig. 1A. The set of simulated FRET time-trajectories were analysed in a similar manner as for the experimentally obtained dataset. Analysis of the simulated dataset (200-ms per frame) showed rates of clamp opening motions, k_closed-partly_closed_, k_partly_closed-open_ and k_closed-open_ to be ∼ 1.3 s^-1^, 0.6 s^-1^ and 0.3 s^-1^, respectively, and, rates of clamp closing motions, k_partly_closed-closed_, k_open-partly_closed_ and k_open-closed_ to be ∼ s^-1^, 1.2 s^-1^ and 0.1 s^-1^, respectively (Fig. S2). We observed three main differences between the simulated dataset and experimental dataset (both with frame rate of 200-ms):

a. The transition rates for the experimental dataset (∼0.02-0.60 s^-1^) are lower than that obtained from the simulated dataset (∼0.10-1.3 s^-1^) by ∼2-5 fold (Fig. S1, S2).
b. A visual inspection of the FRET time trajectories in the simulated dataset (Fig. S2) did not reveal any time trajectories like those on the experimental dataset (Fig. 1C), which show very long-lived clamp conformational states (stable for >20 s).
c. FRET histograms obtained from the simulated dataset differ from that obtained for the dynamic molecules in the experimental dataset, with ∼27%, 54% and 19% of the time spent in the open, partly closed and closed clamp state, respectively, in the simulated dataset as compared to ∼36%, 43% and 21% of the time spent in the open, partly closed and closed clamp state, respectively, in the experimental dataset.

Since the simulations were performed at a frame rate of 200-ms using transition rates from the experimental dataset recorded at a frame rate of 20-ms, the simulated dataset should be able to recapitulate all features which result from missed dwells. A large difference between experimental and simulated dataset, as noted above, indicate that the result of experiments performed at a frame rate of 200 ms cannot be explained solely as an effect of missed dwells. These results strongly suggest that extension of the observation span enables us to capture dwells with hidden transitions between long-lived and short-lived clamp conformational states which are structurally similar in terms of the clamp conformation (i.e., having similar FRET values), and this is reflected in the time trajectories which show very long-lived clamp conformational states (Fig. 1C). Additionally, we note that, from an analysis of the simulated dataset, that in the absence of any hidden transitions, the average dwell times in the respective clamp conformational states for 200-ms frame experiments should be ∼0.8 s,0.8 s and 0.6 s for open, partly closed and closed clamp states. A careful look at the dwells corresponding to the different clamp conformational states obtained from simulations performed at a frame rate of 200-ms show that the number of really long dwells (>5.0 s) corresponding to open clamp (∼0.5%), partly closed clamp (∼5%) and closed clamp (∼0.1%) conformational states were only few in number. In contrast, the experimentally obtained mean dwell times for each state were much longer, for open clamp (∼3.1 s), partly closed clamp (∼2.0 s) and closed clamp (∼1.71 s) conformational states, and the relative number of really long dwells (>5.0 s) were significantly more for open clamp (∼29%), partly closed clamp (∼19%) and closed clamp (∼11%) states. We suggest therefore that these very long dwells in the experimental datasets correspond to either long-lived clamp conformational states or compound dwells consisting of both long-lived and short-lived states with one or more hidden transitions between them.

### The RNAP clamp exhibits unchanged clamp dynamics in the presence of elevated monovalent-cation concentrations

For all our previous experiments on clamp conformational dynamics, we used buffers with 100 mM monovalent (K^+^) and 10 mM divalent (Mg^2+^) salt concentrations. To check if salt with either monovalent or divalent cations influence the clamp status or dynamics, we used increased concentration of each of K^+^ and Mg^2+^ to monitor clamp dynamics, in separate experiments performed as described previously at a frame rate of 20 ms. In a first experiment, we tested the effect of increased concentration of K^+^ (1000 mM) on the status of clamp conformation. Our results showed a similar distribution of clamp states as compared to experiments performed in presence of 100 mM K^+^ (Figure 2A). Specifically, we observed three clamp states: open, E*∼0.18 (43%); partly closed, E*∼0.29 (42%) and closed, E*∼0.49 (15%) (Figure 2A, *left*), with 66% “static” molecules (Figure 2A, *middle*) and 34% “dynamic” molecules (Figure 2A, *right*). Further, estimation of transition rates from the dynamic molecules shows clamp opening rates, k_closed-partly_closed_, k_partly_closed-open_ and k_closed-open_ of ∼ 6.0 s^-1^, 1.0 s^-1^ and 0.1 s^-1^, respectively, and, rate of clamp closing motions, k_partly_closed-closed_, k_open-partly_closed_ and k_open-closed_ of ∼ 2.2 s^-1^, 1.8 s^-1^ and 0.1 s^-1^, respectively (Table 1). Since the overall distribution, the relative proportion of “static” and “dynamic” molecules, and the estimated transition rates are all similar to our results in presence of 100 mM K^+^ (Figure 1A; Table 1), we conclude that an elevated concentration of K^+^ does not have any significant effect on clamp conformation and dynamics.

### The RNAP clamp exhibits decreased clamp dynamics in the presence of elevated divalent-cation concentrations

We also monitored the clamp conformation in presence of increasing concentration of Mg^2+^ (no Mg^2+^, 25 mM Mg^2+^ and 100 mM Mg^2+^). In presence of no Mg^2+^, the equilibrium population distributions indicated the presence of ∼38% of open clamp, ∼38% of partly closed clamp and 24% of closed clamp state, with clamp-opening rates k_closed-partly closed_, k_partly closed-open_ and k_closed-open_ of ∼ 2.8 s^-1^, 0.7 s^-1^ and 0.1 s^-1^, respectively, and clamp-closing rates k_partly closed-closed_, k_open-partly closed_ and k_open-closed_ of ∼ 0.7 s^-1^, 0.7 s^-1^ and 0.05 s^-1^, respectively; these results were similar to those obtained for clamp conformation and dynamics in presence of 10 mM Mg^2+^ (no Mg^2+^, Figure 2B, *first row*; and 10 mM Mg^2+^, Figure 1A, *left* and Table 1). However, a significant change in the clamp conformation distribution was observed at 25 mM Mg^2+^, where our data fit best to a two-state model with clamp populating only open (E*∼0.22; 69%) and closed (E*∼0.41; 31%) conformations. Surprisingly, upon increasing the Mg^2+^ concentration to 100 mM, we observed a dramatic shift in the clamp conformation distribution to almost an exclusively open (E*∼0.20; 91%) conformation (Figure 2B, *left panels*) and a significant reduction in “dynamic” molecules (∼11% for 25 mM Mg^2+^; ∼9% for 100 mM Mg^2+^; Figure 2B, *right panels*), with most molecules exhibiting “static” clamp status (∼89% for 25 mM Mg^2+^; ∼91% for 100 mM Mg^2+^; Figure 2B, *middle panels*). We conclude that elevated concentrations of Mg^2+^ shift the conformational equilibria to a long-lived open clamp conformation.

### The RNAP clamp exhibits increased clamp dynamics in the presence of molecular crowding

In the context of a living cell, the RNAP exists in a complex mixture of other proteins, nucleic acids and small molecules. It has been estimated that the overall cellular milieu consists of ∼80-400 mg/ml of other macromolecules which occupy up to ∼30% of volume inside the cell (*20,21*). Such crowded environments have been reported to exert an “excluded-volume” effect resulting in a shift towards more compact conformations of the protein (*22,23*). Most *in-vitro* studies looking at protein behaviour in crowded environments tend to use “inert crowders” like polyethylene glycol (PEG) or dextran which mimic the excluded-volume effect (*22*).

To examine how a crowded cellular milieu may affect RNAP clamp conformational dynamics, we used 20% polyethylene glycol (PEG-8000) in the imaging buffer to study motions of clamp at a frame rate of 20-ms. Our results show that, in presence of 20% PEG, the RNAP clamp populates three conformational states: open (E*∼0.19; 28%), partly closed (E*∼0.32; 58%) and closed (E*∼0.56; 14%) (Figure 3A, *left*). These results show an overall depopulation of the open clamp state and increase in relative abundance of the partly closed clamp state (Figure 3A, *top right*) compared to the non-crowded conditions. Additionally, we observe a shift in FRET efficiencies for the partly closed and closed clamp conformation to higher FRET values (0.32 compared to 0.29 for the partly closed, and 0.56 compared to 0.42 for the closed clamp state), indicative of a greater degree of clamp closure in these states. Both observations are consistent with expectations that macro-molecular crowding may result in stabilisation of more compact protein conformations (*24,25*), and in this case, greater degree of clamp closure within RNAP molecules. HMM analysis of time trajectories for this dataset reveals only ∼43% of the RNAP molecules were “static” (compared to ∼70% molecules for the non-crowded condition; Figure 3A, *top left*), whereas most molecules showed a dynamic behaviour (∼57%; Figure 3A, *bottom left*). Most notably, estimates of transition rates for clamp-opening motions, k_closed-partly_closed_ and k_partly_closed-open_ were ∼ 7.7 s^-1^, and 1.2 s^-1^, respectively, and, rate of clamp closing motions, k_partly_closed-closed_ and k_open-partly_closed_ were ∼ 1.7 s^-1^ and 2.5 s^-1^, respectively (Fig. 3A, *bottom right*; Table 1), under the conditions of crowding, indicating ∼2-fold enhancement in rates of switching between the different short-lived clamp conformations. Direct transitions between open and closed conformations were rarely observed and as such it was not possible to estimate transition rates between them. Taken together, the increased proportion of “dynamic” molecules and the increased transition rates indicate an overall enhancement of clamp motions under conditions of macromolecular crowding.

### The RNAP clamp predominantly exhibits a closed conformation when RNAP is bound non-specifically to DNA

It has been proposed that RNAP can bind to non-specific DNA in the cell as it tries to find a promoter sequence via either 1D-sliding along genomic DNA, or via a 3D-diffusion-mediated “hop and search” mechanism, or a combination of both (*29-31*). Further, it has been proposed that, while searching for promoters along the bacterial genome, the clamp populates an open conformation and undergoes transient clamp closures as RNAP tries to locate a promoter (*5*). Such proposals however, have been speculative, as no direct observation of clamp conformations or motions during promoter search has been made to date. To gain insight into RNAP clamp conformation and dynamics at non-specific DNA binding sites as would be encountered by the RNAP while searching for promoter sites, we designed a biotinylated non-specific DNA by changing the segments (−35 and −10 positions) of a *lac*-promoter responsible for the sequence-specific recognition by RNAP (see Methods). In a next set of experiments, we pre-formed complexes with RNAP and biotinylated non-specific DNA, immobilised pre-formed single RNAP-non-specific DNA complexes on a PEGylated glass surface via biotin on DNA, and recorded single-molecule FRET time-trajectories at frame rates of 100 ms. Since immobilisation was performed via biotin on DNA, all RNAP molecules under observation were bound to non-specific DNA; in fact, control experiments showed that under our conditions, no RNAP molecules stay bound to the surface in a non-specific manner (i.e., in the absence of DNA). Relative abundances of clamp conformational states for all RNAP molecules analysed were 23%, 24% and 53% for open, partly closed and closed clamp conformations, respectively (Figure 3B, *top right*; histogram from all molecules). HMM analysis of the trajectories for single RNAP-non-specific-DNA complexes revealed that the clamp populates a “static” conformation for the majority (∼85%) of molecules (Figure 3B, *top left*) and show “dynamic” behaviour for ∼15% of the molecules (Figure 3B, *bottom left*) with estimated rate of clamp opening motions, k_closed-partly_closed_ and k_partly_closed-open_ of ∼ 0.5 s^-1^, and 0.3 s^-1^, respectively, and, rate of clamp closing motions, k_partly_closed-closed_ and k_open-partly_closed_ of ∼ 0.3 s^-1^ and 0.1 s^-1^, respectively (Fig. 3B, *bottom right*; Table 1). Direct transitions between the open and closed clamp conformations were rare. We conclude that, when bound to non-specific DNA, the clamp exists in a predominantly closed conformation, but can also show very slow opening-closing dynamics. This is in sharp contrast with results obtained in similar experiments with preformed RNAP-promoter complexes, where the clamp was found to be in a stably closed status and did not exhibit any dynamic behaviour (*13,15*).

## Discussion

In this work, we build on our previous smFRET studies monitoring RNAP clamp conformation and conformational dynamics in solution (*13, 15, 16*) by extending the observation span of our experiments to capture structurally similar but kinetically different long-lived clamp conformations and by monitoring the clamp conformation and motions under environments RNAP molecules may encounter in a living cell, such as molecular crowding, elevated monovalent and divalent cation concentrations, and non-specific DNA binding.

Our previous smFRET studies revealed two classes of RNAP molecules depending on the nature of clamp motions: “static” molecules, which show no transition to other clamp conformations, and “dynamic” molecules, which show interconversion between the different clamp conformations. Here, using an extended observation span (∼120 s), we show that previously observed molecules with a “static” clamp conformation are not intrinsically different to the “dynamic” molecules, but actually result from the presence of structurally similar but kinetically distinct long-lived clamp conformational sub-states, as a result of which several molecules appear to be “static” at a particular clamp conformation during short observation span (∼10 s) experiments. With the help of simulations and detailed analysis of new experiments performed at decreased frame-times (∼200 ms), we uncover the existence of long-lived clamp conformational sub-states that last for >1.0 s, a timescale at least ∼3-fold longer than average dwell times in the short-lived conformational sub-states of ∼0.3-0.6 s. Inspection of individual time-trajectories shows clear evidence of interconversion between these newly observed long-lived and previously observed short-lived clamp conformational sub-states. We note that the transition rates between these conformational sub-states potentially could result in an additional, previously unknown, point for modulation of clamp dynamics.

### Modulation of clamp dynamics by monovalent and divalent cation concentrations

We have also shown that, intriguingly, high concentration (>25 mM) of a salt with a divalent cation (Mg^2+^) shifts the conformational equilibria to an open clamp state, essentially abrogating any clamp-closing motions. This observation provides a simple chemical means to stabilise the open clamp conformation for further study. On the other hand, the biological relevance of the Mg^2+^ dependence is unclear, since the concentration of free Mg^2+^ inside bacterial cells is only ∼ 1 mM (*33*). However, a much large amount of cellular Mg^2+^ (∼20-100 mM) is bound to other macromolecules (ribosomes, ribozymes, nucleic acids) (*33-35*), and under certain cellular conditions, e.g., stress induced by antibiotics targeting ribosomes may result in higher magnesium flux in a bacterial cell (*36*). Such a condition may result in a Mg^2+^-dependent trapping of the RNAP clamp in an open conformation, resulting in down-regulation of transcription.

### Modulation of clamp dynamics by molecular crowding

The bacterial cell also presents a crowded environment for proteins and can dramatically affect thermodynamic equilibria, diffusion properties and binding between interacting partners due to excluded-volume effects. Our results using a “macro-molecular crowder” (20% PEG-8000), which is expected to mimic conditions of a crowded cellular milieu (*22*), show a shift to a more closed conformation for the partly closed and closed clamp conformations, and an increase in the relative abundance of the partly closed state (Figure 2A) when compared with non-crowded conditions. This is consistent with the expectation that macromolecular crowding stabilises compact conformations of a protein (*24*). More significantly, under crowded conditions, the number of static molecules (∼43%) were significantly lower than that observed under non-crowded conditions (∼70%), and the estimated clamp opening and clamp closing rates are ∼2-fold higher compared to experiments under non-crowded conditions. These results suggest that, in the crowded cellular environment, intrinsic conformational dynamics of the clamp are significantly enhanced, possibly via a relative stabilisation of the transition states (Fig. 4). We thus suggest that changes in intracellular crowding (e.g., micro-compartmentalisation due to possible liquid-liquid phase separation; *28*) may further alter clamp conformational dynamics and modulate the overall rate of transcription initiation. Future studies monitoring conformational dynamics of RNAP clamp in live bacterial cells would be important to understand such aspects of transcription regulation.

**Fig. 4.**
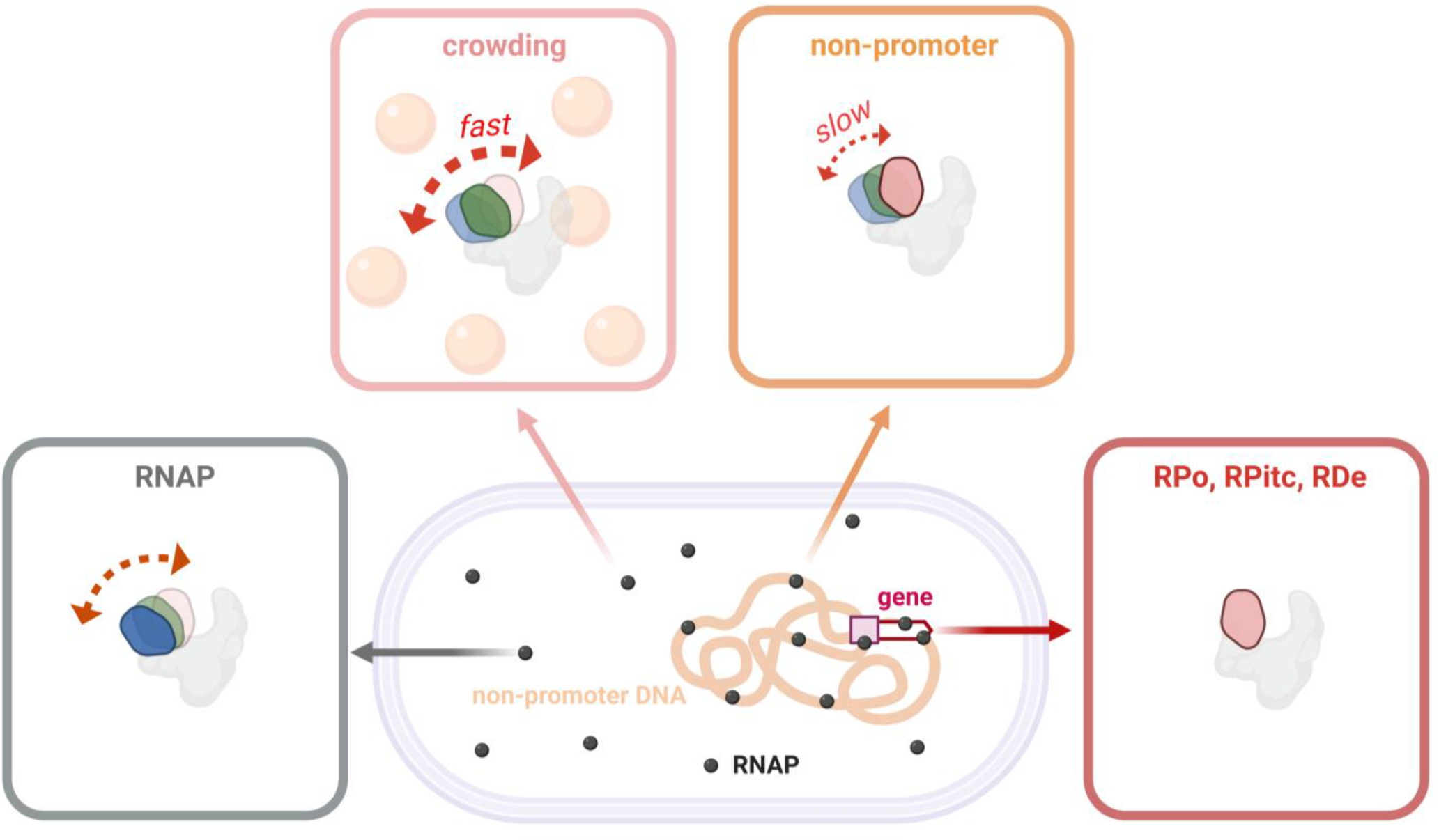
Summary of clamp conformational status in free and bound RNAP. Schematic showing clamp conformation and dynamics in free RNAP, RNAP in presence of macromolecular crowding, RNAP complexes bound to non-specific DNA, and RNAP during different stages of transcription. Black dots, RNAP; blue, open clamp; green, partly closed clamp; red, closed clamp; grey, rest of RNAP; double headed dotted arrows, clamp motions; light orange spheres, macromolecular crowders; orange, bacterial nucleoid; rectangular purple box, promoter; white arrow, gene.

### Modulation of clamp dynamics during promoter search

Transcription initiation is one of the major targets of gene regulation; part of the regulation may occur via modulation of the process by which RNAP “scans” the bacterial nucleoid to locate and bind to promoter DNA, a process also known as “promoter search”. Promoter search has been proposed to occur via either a 3D-diffusion mediated “hop and search” mechanism involving transient binding-unbinding to non-specific DNA, via 1D-diffusion along the bacterial nucleoid involving sliding along non-specific DNA until it encounters a promoter sequence, or via a combination of these processes (*29-31, 36*). Recent studies tracking single RNAP molecules in single live bacterial cells show that RNAP spends the large majority of its search time (∼85%) engaged non-specifically with bacterial nucleoid (*32*).

It has been hypothesized that the RNAP holoenzyme mainly adopts an open-clamp conformational state, and only transiently samples closed-clamp conformational states, when RNAP holoenzyme interacts with non-specific DNA during promoter search (*5*). Our results argue against this hypothesis. We find that RNAP holoenzyme mainly adopts a *closed-clamp* conformational state--with slow transitions between closed, partly closed, and open states--when RNAP holoenzyme interacts with non-specific DNA (Fig. 4). However, although our results argue against the proposal of a mostly open clamp RNAP molecule sliding along the genomic DNA with transient clamp closures punctuating the promoter search process, the observation that some slow clamp opening-closing transitions do take place when bound to non-specific DNA, raises the possibility that clamp conformational dynamics may play a role during promoter search process. Recent studies combining single-molecule fluorescence measurements with optical tweezers have been able to observe long range 1D-sliding of RNAP molecules on doubly tethered λ DNA (*37*). A combination of methods used in the aforementioned study with reagents developed here should enable us to capture clamp conformations of actively diffusing RNAP molecules along DNA and provide more mechanistic detail about the role of clamp dynamics during facilitated diffusion of RNAP.

## ACKNOWLEDGEMENTS

We thank Wellcome Trust [110164/Z/15/Z to A.N.K.]; NIH [GM041376 to R.H.E.]; Funding for open access charge: Wellcome Trust [110164/Z/15/Z], for funding this work.

## Author Contributions

A.N.K., R.H.E., and A.M conceived the project; A.M. prepared and characterized protein samples; A.M. and A.W. performed single-molecule experiments; A.M. and A.W. analyzed data; A.W. and H.U. performed simulations; A.M. and A.W. prepared figures; and A.M., R.H.E., and A.N.K wrote the manuscript.

## Supplementary Figures

**Fig. S1:**
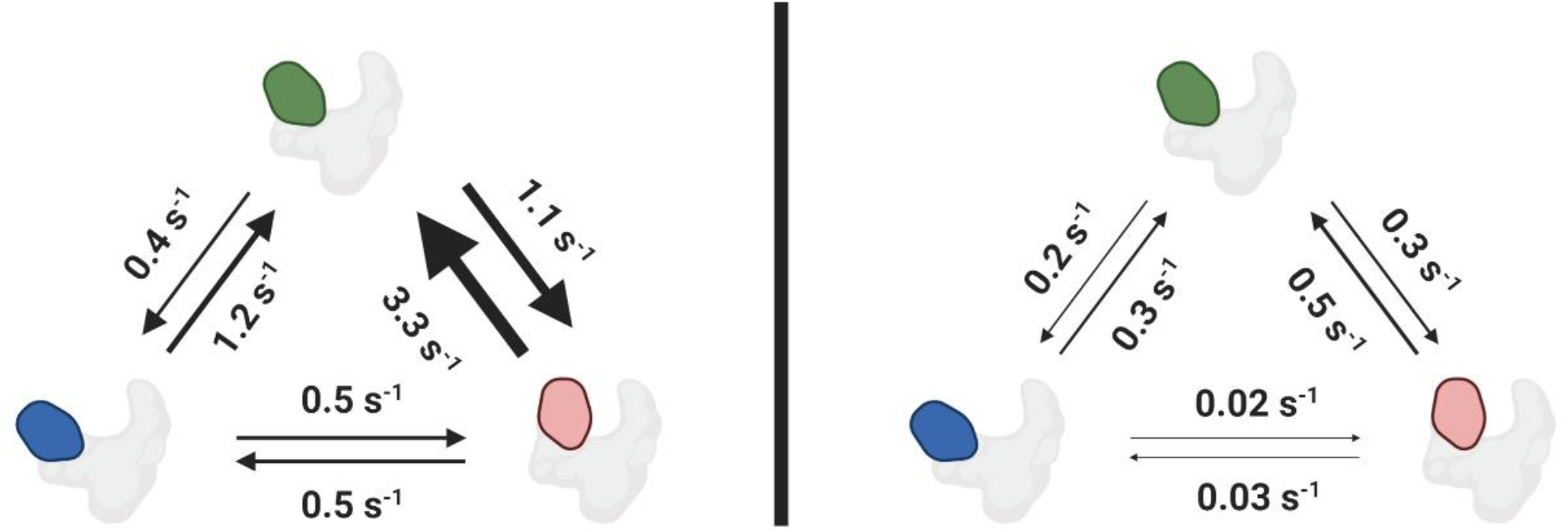
Transition rates between open, partly closed and closed conformations obtained from experiments performed at a frame rate of 20-ms (*left*) and 200-ms (*right*). Blue, open clamp; green, partly closed clamp; red, closed clamp; grey, rest of RNAP.

**Fig. S2.**
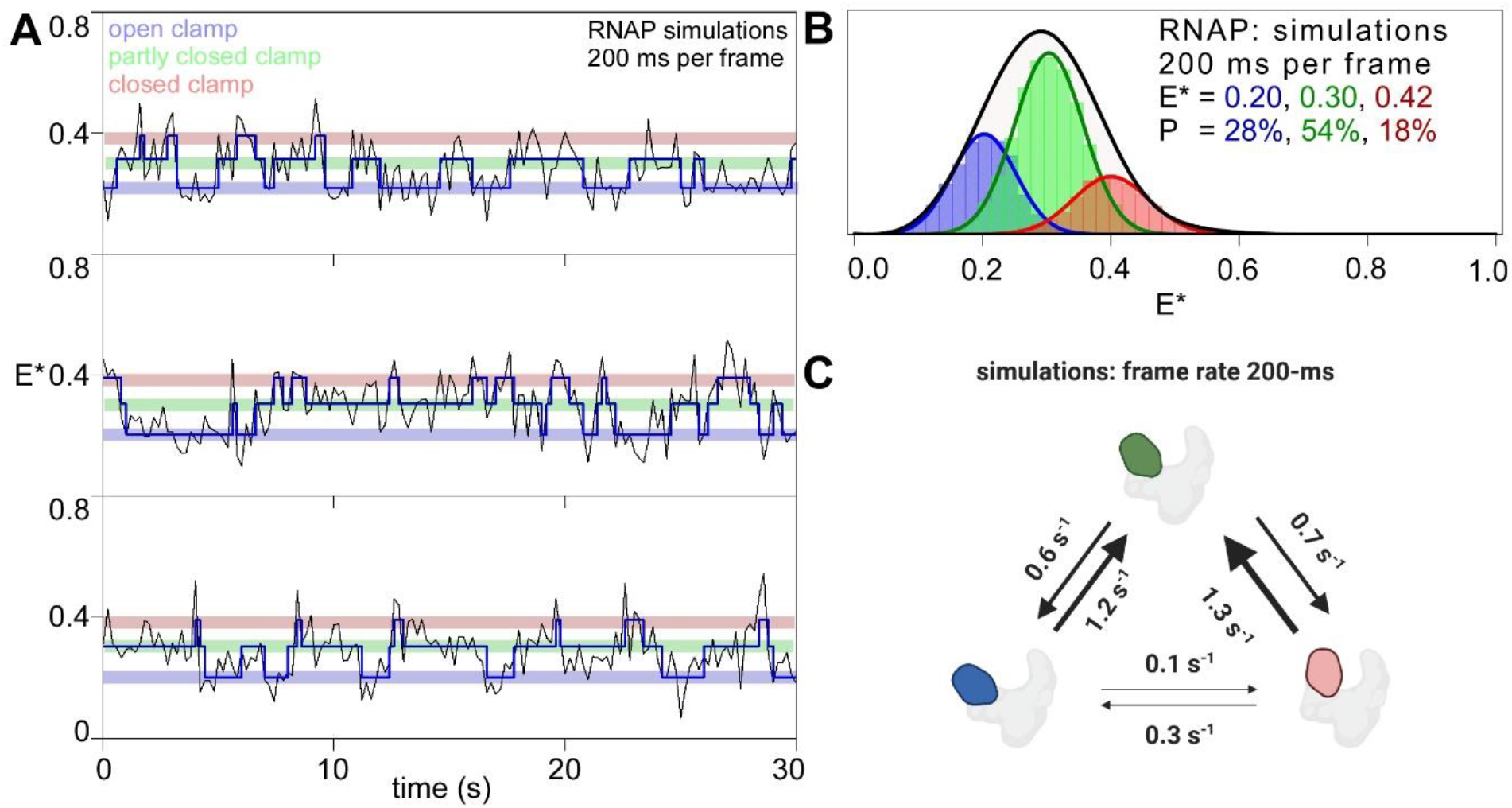
**A.** Representative time traces of FRET efficiency, E*, showing hidden-Markov-model (HMM)-assigned states for simulations performed at a frame rate of 200-ms using transition rates obtained from experiments with frame rate of 20-ms; open (faded blue), partly closed (faded green), closed clamp (faded red) states, and interstate transitions (blue line). **B**. Histograms and Gaussian fits of E*, showing HMM-derived E* distributions for simulations performed at a frame rate of 200-ms using transition rates obtained from experiments with frame rate of 20-ms; open (blue bars), partly closed (green bars) and closed (red bars) clamp states; P, subpopulation percentage. **C**. Transition rates between open, partly closed and closed conformations obtained from simulations performed at a frame rate of 200-ms. Blue, open clamp; green, partly closed clamp; red, closed clamp; grey, rest of RNAP.

